## Supplement for "Assessment of false discovery rate control in tandem mass spectrometry analysis using entrapment"

### S1 An abstraction of the entrapment concept

This section lays out a proposed abstract formulation of the entrapment procedure. We consider an analysis tool that takes an input dataset $\mathcal{I}$ and produces a list of putative discoveries $\{\delta_i\}$. The discoveries are ranked by their scores $\{S_i\}$ that summarize the evidence in support of the discoveries, so that the larger $S_i$ is, the more likely $\delta_i$ is to be a true or correct discovery. For example, in the MS/MS context $\mathcal{I}$ can consist of the set of MS2 spectra and the target peptide database. A PSM-level analysis tool in this context can return a ranked list of scored target PSMs, where an incorrect match corresponds to a false discovery. Similarly, a peptide-level analysis tool can return a list of target peptides with at least one matching PSM, along with the score of the maximal PSM associated with the peptide. In this case, any reported peptide that is missing from the sample is a false discovery. Note that a precursor-level analysis is conceptually the same as the peptide-level one, except that the considered objects are specifically-charged (possibly modified) peptides rather than just peptides, so below we do not distinguish between the two.

Furthermore, given an FDR threshold $\alpha$, the tool reports a subset of its putative discoveries while aiming to control the FDR among those reported. To simplify the exposition we assume here that the selected list of discoveries is made, as is typically the case, of the top $K_\alpha$ ranked ones, but our arguments apply more generally. The expectation with respect to which the FDR is defined is taken over any randomness in the generation of the data $\mathcal{I}$, as well as any that is built into the tool (e.g., if it randomly draws decoys).

The goal of the entrapment procedure is to gauge whether the analysis tool, which could be a black box, correctly controls the FDR in its reported lists of discoveries. To do that, the entrapment procedure randomly expands $\mathcal{I}$ to a larger dataset $\bar{\mathcal{I}}$ designed to force the tool to consider verifiably false putative discoveries. For example, in the MS/MS context we typically expand the original target database by adding to it entrapment peptides that we believe are not present in the sample. Critically, the expansion needs to be done so that entrapment discoveries realistically model false original discoveries. Subsequently, the procedure applies the analysis tool to the entrapment-expanded $\bar{\mathcal{I}}$ to derive a combined list of original and entrapment discoveries.

At the last step, the entrapment procedure employs one or more estimation methods—some of which are explored in the next section—to estimate the FDP in that combined list of discoveries. By assumption, any entrapment discovery is a false one, but an estimation method can leverage those entrapment discoveries to estimate the number of false original discoveries. Note that, as defined here, the sample estimation approach, Equation (3), does not qualify as an entrapment procedure because it does not aim to estimate the FDP in the combined list.

### S2 Estimation methods and the assumptions they rely on

| Name | Notation | Expression | Required Assumptions and Comments |
|---|---|---|---|
| Lower Bound | $\underline{\widehat{\text{FDP}}}_{\mathcal{E}\,\cup\,\mathcal{O}}$ | $\frac{N_{\mathcal{E}}}{N_{\mathcal{O}}+N_{\mathcal{E}}}$ | Every entrapment discovery is a false one (Assumption 1, see also Assumptions 1a and 1b for common use cases). |
| Combined (average upper bound) | $\widehat{\text{FDP}}_{\mathcal{E}\,\cup\,\mathcal{O}}$ | $\frac{N_{\mathcal{E}}(1+1/r)}{N_{\mathcal{O}}+N_{\mathcal{E}}}$ | Given the number of discoveries $K_\alpha$, any false discovery is at least $r$ times more likely to be an entrapment than an original discovery (Assumption 2). See also Assumption 2a and the discussion following it for common use cases. |
| Matched (average upper bound) | $\widehat{\text{FDP}}^{*k}_{\mathcal{E}\,\cup\,\mathcal{O}}$ | $\frac{N_{\mathcal{E}}+\sum_{l=1}^{k+1} l\cdot N_l}{N_{\mathcal{O}}+N_{\mathcal{E}}}$ | Every potential original discovery is matched with $k$ potential entrapment discoveries. Roughly, when ordered together with its $k$ matched entrapments a potential original false discovery is equally likely to occupy each of the $k+1$ possible ranks independently of how many of these $k+1$ are reported discoveries and of the total number of discoveries $K_\alpha$(Assumption 3). See also Assumption 3a and the discussion following it for common use cases. For $k=1$ this is the paired estimation, Equation (4). |

Table S1: **Summary of entrapment estimation methods discussed in this section.** Note that the required assumptions for the upper bound methods are variants of the equal chance assumption that TDC relies on: an incorrect discovery is equally likely to be a decoy or a target. Notably, the entrapment estimation does not require independence between the false discoveries, which TDC requires and which makes PSM-level FDR control particularly challenging.

#### S2.1 The lower-bound estimation

We start with some required notation:

- $N_{\mathcal{O}}$ and $N_{\mathcal{E}}$ denote the number of original and entrapment discoveries, respectively, that are reported when analyzing the expanded dataset $\bar{\mathcal{I}}$ at the given FDR threshold $\alpha$.
- $K_\alpha$ denotes the number of top discoveries reported by the analysis tool at the given FDR threshold $\alpha$, so that $N_{\mathcal{O}} + N_{\mathcal{E}} = K_\alpha$.

The lower bound estimation method is defined analogously to Equation (2) with $N_{\mathcal{O}}$ (number of original discoveries) replacing the MS/MS specific $N_{\mathcal{T}}$ (number of target discoveries):

$$\underline{\widehat{\text{FDP}}}_{\mathcal{E}\,\cup\,\mathcal{O}} = \frac{N_{\mathcal{E}}}{N_{\mathcal{O}} + N_{\mathcal{E}}}. \tag{S1}$$

The lower bound relies on

**Assumption 1.** *Every entrapment discovery is a false one.*

In the context of MS/MS entrapment through database expansion, we can satisfy this assumption for the PSM-level, and peptide-level analyses through

**Assumption 1a.** *None of the entrapment peptides is present in the sample.*

Similarly, for protein-level analysis we can satisfy Assumption 1 through

**Assumption 1b.** *None of the entrapment proteins is present in the sample.*

**Claim 1.** If Assumption 1 holds then $\underline{\widehat{\text{FDP}}}_{\mathcal{E}\,\cup\,\mathcal{O}}$ is a rigorous lower bound on the FDP among the combined list of $N_{\mathcal{E}} + N_{\mathcal{O}}$ entrapment and original discoveries.

Indeed, this is a trivial observation: let $\mathring{N}_{\mathcal{O}}$ denote the number of false discoveries among the $N_{\mathcal{O}}$ reported original discoveries. Then under Assumption 1, the actual FDP among the reported discoveries is given by $(\mathring{N}_{\mathcal{O}} + N_{\mathcal{E}})/K_\alpha \geq N_{\mathcal{E}}/K_\alpha$.

#### S2.2 The combined estimation

The combined method involves a parameter $r$ whose significance is explained in the next assumption. However, that assumption first requires the following notation:

- $\Phi$ denotes the set of reported false discoveries (original or entrapment), and
- $\mathcal{O}$ and $\mathcal{E}$ denote the sets of reported original and entrapment discoveries, respectively.

**Assumption 2.**

$$P(\delta \in \mathcal{E} \,|\, \delta \in \Phi \,,\, K_\alpha) \geq r \cdot P(\delta \in \mathcal{O} \,|\, \delta \in \Phi \,,\, K_\alpha).$$

*The probability is taken with respect to the random effects in the generation of the original data $\mathcal{I}$, its random expansion to $\bar{\mathcal{I}}$ by the entrapment procedure, as well as any random effects that the analysis tool might introduce.*

In words, given the number of discoveries $K_\alpha$, any false discovery is at least $r$ times more likely to be an entrapment than an original discovery. Going back to the context of MS/MS entrapment through database expansion, we can, for example, satisfy Assumption 2 in PSM-, peptide- and protein-level analyses by assuming that the expansion guarantees the following:

**Assumption 2a.** *An incorrect PSM, peptide or protein is at least $r$ times more likely to be an entrapment than a target one, independently of the number of discoveries.*

In practice, there is a difference in how the last assumption is interpreted at the PSM level compared with the peptide and protein level. Specifically, at the PSM level we typically assume that if the ratio of entrapment to target peptides is $r : 1$ then an incorrect PSM is (exactly) $r$ times more likely to be an entrapment than a target one. Indeed, for $r = 1$ this is analogous to the equal chance assumption that TDC requires for the relation between decoys and targets. Moreover, this assumption is reasonable here if we assume that (a) the entrapment peptides realistically model peptides that are not in the sample, and (b) because the search tool has no access to the target/entrapment labels, this equal chance happens regardless of how many discoveries the tool reports.

For peptide- and protein-level analyses the "at least" part of Assumption 2a becomes critical. Indeed, in these cases a false discovery corresponds to the reported protein or peptide missing from the sample. Therefore, considering for simplicity the case $r = 1$, even when there is a 1:1 correspondence between target and entrapment peptides, and even if the latter adequately model target peptides that are missing from the sample, then given that a peptide is missing it is more likely to be an entrapment than a target: some of the target peptides are present in the sample, whereas none of the entrapment ones should be. So in this case a false discovery should be more likely to be an entrapment. The more general assumption corresponds to when the ratio of entrapment peptides (proteins) to target ones is $r : 1$, again assuming that (a) the entrapment peptides (proteins) realistically model those that are not in the sample, and (b) the search tool has no access to the target/entrapment labels.

The combined estimation method relies on Assumption 2 to give an (averaged) upper bound on the FDP, defined analogously to Equation (1):

$$\widehat{\text{FDP}}_{\mathcal{E} \,\cup\, \mathcal{O}} = \frac{N_{\mathcal{E}}(1 + 1/r)}{N_{\mathcal{O}} + N_{\mathcal{E}}}. \tag{S2}$$

The precise, rigorous nature of this upper bound is described next.

**Claim 2.** If Assumption 2 holds then the expected value of $\widehat{\text{FDP}}_{\mathcal{E} \,\cup\, \mathcal{O}}$ provides a rigorous upper bound on the FDR among the combined list of $K_\alpha = N_{\mathcal{E}} + N_{\mathcal{O}}$ entrapment and original discoveries:

$$E(\widehat{\text{FDP}}_{\mathcal{E} \,\cup\, \mathcal{O}}) \geq E\left[\frac{N_{\mathcal{E}} + \mathring{N}_{\mathcal{O}}}{K_\alpha}\right],$$

where the expectations are taken with respect to the same random factors as in Assumption 2.

Note that while $\underline{\widehat{\text{FDP}}}_{\mathcal{E} \cup \mathcal{O}}$ provides a lower bound on the FDP (and hence, if averaged, also on the FDR), the combined estimate $\widehat{\text{FDP}}_{\mathcal{E} \cup \mathcal{O}}$ provides an indirect upper bound on the FDP: its expected value bounds the expected value of the FDP among the reported discoveries, i.e., the FDR. It is also worth highlighting that with $r = 1$, Assumption 2 is similar to the equal chance assumption that TDC relies on. Notably, TDC also requires the independence of the false discoveries, whereas the combined estimation does not.

*Proof.* It follows from Assumption 2 that with $\delta_i$ being the $i$th discovery

$$
\begin{aligned}
E(N_{\mathcal{E}} \mid K_\alpha) &= E(\sum_{i=1}^{K_\alpha} 1_{\delta_i \in \mathcal{E}} \mid K_\alpha) = \sum_{1}^{K_\alpha} E(1_{\delta_i \in \mathcal{E}} \mid K_\alpha) = \sum_{1}^{K_\alpha} P(\delta_i \in \mathcal{E} \mid K_\alpha) \geq \sum_{1}^{K_\alpha} P(\delta_i \in \Phi \, , \, \delta_i \in \mathcal{E} \mid K_\alpha) \\
&= \sum_{1}^{K_\alpha} P(\delta_i \in \mathcal{E} \mid \delta_i \in \Phi \, , \, K_\alpha) \cdot P(\delta_i \in \Phi \mid K_\alpha) \geq \sum_{1}^{K_\alpha} r \cdot P(\delta_i \in \mathcal{O} \mid \delta_i \in \Phi \, , \, K_\alpha) \cdot P(\delta_i \in \Phi \mid K_\alpha) \\
&= r \cdot \sum_{1}^{K_\alpha} P(\delta_i \in \Phi \, , \, \delta_i \in \mathcal{O} \mid K_\alpha) = r \cdot \sum_{1}^{K_\alpha} E(1_{\delta_i \in \mathcal{O} \cap \Phi} \mid K_\alpha) = r \cdot E(\sum_{i=1}^{K_\alpha} 1_{\delta_i \in \mathcal{O} \cap \Phi} \mid K_\alpha) \\
&= r \cdot E(\mathring{N}_{\mathcal{O}} \mid K_\alpha).
\end{aligned}
$$

Therefore,

$$
E\left[ \frac{N_{\mathcal{E}}(1 + 1/r)}{K_\alpha} \mid K_\alpha \right] = \frac{E\left[ N_{\mathcal{E}}(1 + 1/r) \mid K_\alpha \right]}{K_\alpha} \geq \frac{E(N_{\mathcal{E}} \mid K_\alpha) + E(\mathring{N}_{\mathcal{O}} \mid K_\alpha)}{K_\alpha} = E\left[ \frac{N_{\mathcal{E}} + \mathring{N}_{\mathcal{O}}}{K_\alpha} \mid K_\alpha \right].
$$

Taking expectation of both sides of the last inequality completes the proof. □

#### S2.3 The paired and matched estimations

The combined estimation provides a valid average upper bound on the FDP; however, it can be overly conservative, i.e., unduly over-estimate the FDP. Indeed, in peptide-level analysis there can be a significant gap between the factor $r$, denoting the relative size of the entrapment database, and the effective ratio of $P(\delta \in \mathcal{E} \mid \delta \in \Phi \, , \, K_\alpha)/P(\delta \in \mathcal{O} \mid \delta \in \Phi \, , \, K_\alpha)$. Specifically, the discussion following Assumption 2a implies that this latter ratio should in fact be taken as $r/\pi_0$, where $\pi_0$ is the fraction of target database peptides that is missing from the sample. Thus, assuming $\pi_0$ is known, and using essentially the same proof of Claim 2, we can show that $N_{\mathcal{E}}(1 + \pi_0/r)/(N_{\mathcal{O}} + N_{\mathcal{E}})$ is also a valid average upper bound and one which is clearly smaller than $\widehat{\text{FDP}}_{\mathcal{E} \cup \mathcal{O}} = N_{\mathcal{E}}(1 + 1/r)/(N_{\mathcal{O}} + N_{\mathcal{E}})$.

Of course, $\pi_0$ is unknown, but this observation motivated our introduction of tighter average upper bound estimations, the paired method, and its generalization, the matched method. The applicability of the paired method hinges on the data expansion process pairing each potential original discovery with an entrapment one. That is, the expansion is done in such a way that there is a 1:1 and onto mapping $\rho$ that links each potential original discovery to a unique potential entrapment discovery and vice versa. For example, we can achieve this in the context of peptide-level analysis if we generate $\bar{\mathcal{I}}$ by expanding the target database by pairing each target database peptide with a unique, shuffled entrapment peptide.

The matched method relies on a generalization of this pairing by assuming that the expansion is done in such a way that each potential original discovery is matched to $k$ distinct potential entrapment discoveries. For example, in the peptide-level analysis context we would generate $\bar{\mathcal{I}}$ by expanding the target database through matching each target database peptide with $k$ distinct shuffled entrapment peptides.

To introduce and analyze the matched method we need the following notation:

- $\Delta = \mathcal{O} \cup \mathcal{E}$ denotes the set of discoveries.
- $\bar{\mathcal{O}}$ denotes the set of all original potential discoveries (e.g., all target database peptides in the peptide-level analysis).
- $\bar{\Phi}$ denotes the set of all potential false discoveries (original or entrapment),

- For $\delta \in \bar{\mathcal{O}}$, $\Pi_\delta$ denotes the set of $k+1$ potential discoveries that include $\delta$ as well as its $k$ matched entrapments (e.g., for the paired method $\Pi_\delta = \{\delta, \rho(\delta)\}$).
- $R_\delta$ denotes the rank of $\delta \in \bar{\mathcal{O}}$ among the scores of the $k+1$ potential discoveries in their $\Pi_\delta$, where $R_\delta = k+1$ when $\delta$ is ranked last. Any potential entrapment or original discovery that is not scored by the tool is necessarily absent from $\Delta$ as well, and it is given a score of $-\infty$. Ties are broken randomly.
- $N_l$ denotes the number of original potential discoveries $\delta \in \bar{\mathcal{O}}$, which are ranked last in $\Pi_\delta$, and for which exactly $l$ of the potential discoveries in their $\Pi_\delta$ made it into $\Delta$:

$$N_l = \left|\left\{\delta \in \bar{\mathcal{O}} \,:\, \#\left(\Pi_\delta \cap \Delta\right) = l \,,\, R_\delta = k+1\right\}\right|.$$

- $\mathring{N}_l$ denotes the number of original false discoveries $\delta$ for which exactly $l$ of the potential discoveries in their $\Pi_\delta$ made it into $\Delta$:

$$\mathring{N}_l = \left|\left\{\delta \in \mathcal{O} \cap \Phi \,:\, \#\left(\Pi_\delta \cap \Delta\right) = l\right\}\right|.$$

The $k$-matched estimation method is defined as

$$\widehat{\text{FDP}}^{*k}_{\mathcal{E} \cup \mathcal{O}} = \frac{N_{\mathcal{E}} + \sum_{l=1}^{k+1} l \cdot N_l}{N_{\mathcal{O}} + N_{\mathcal{E}}}. \tag{S3}$$

Note that for $k = 1$ this coincides with the paired method and particularly with Equation (4) that we introduced in the context of peptide-level analysis.

**Assumption 3.** *For $j, l \in \{1, 2, \ldots, k+1\}$*

$$P\left(R_\delta = j \,\middle|\, \delta \in \bar{\mathcal{O}} \cap \bar{\Phi} \,,\, \#\left(\Pi_\delta \cap \Delta\right) = l \,,\, K_\alpha\right) = \frac{1}{k+1},$$

*where the probability is defined as in Assumption 2.*

In words, considering a potential original false discovery $\delta \in \bar{\mathcal{O}} \cap \bar{\Phi}$ (e.g., a target peptide that is not in the sample), given that the set $\Pi_\delta$ (which contains $\delta$ and its $k$ matched entrapments) has exactly $l$ discoveries, and given the overall number of discoveries $K_\alpha$, the score of $\delta$ is drawn from the same distribution as that of its matched entrapment: its rank in the combined set $\Pi_\delta$ is equally likely to be any of the $k+1$ possible ranks. Returning to the context of MS/MS entrapment through database expansion, we can for example satisfy Assumption 3 in peptide- and protein-level analyses by assuming that the expansion guarantees the following:

**Assumption 3a.** *The score of a target peptide that is missing from the sample is drawn from the same distribution of its $k$ matched entrapment peptides, independently of how many discoveries are made overall and of how many discoveries are among the $k+1$ considered peptides (replace "peptide" with "protein" for protein-level analysis).*

In the case $k = 1$ (paired method), this is a reasonable assumption as long as each entrapment peptide offers a realistic alternative to its paired target peptide when the latter is missing from the sample. Indeed, in that case, each one of the two is equally likely to score higher, and because the search engine cannot see the target/entrapment label this should happen independently of whether both peptides or just one of them was discovered, and of how many discoveries were reported overall. This is similar to the equal chance assumption that TDC relies on, and it is worth noting that, like the lower bound and combined methods, the paired estimation does not require the independence of the false discoveries, which TDC does.

Note that there is no obvious way to apply the paired method to PSM-level analysis, because it is not clear how to pair an entrapment PSM with a target PSM. Specifically, pairing peptides does not immediately translate to pairing PSMs unless we already selected the best PSM for each peptide, in which case this is again peptide-level analysis.

**Claim 3.** If Assumption 3 holds then the expected value of $\widehat{\mathrm{FDP}}^{*k}_{\mathcal{E}\,\cup\,\mathcal{O}}$ provides a rigorous upper bound on the FDR among the combined list of $K_\alpha = N_\mathcal{E} + N_\mathcal{O}$ entrapment and original discoveries:

$$E(\widehat{\mathrm{FDP}}^{*k}_{\mathcal{E}\,\cup\,\mathcal{O}}) \geq E\left[\frac{N_\mathcal{E} + \mathring{N}_\mathcal{O}}{K_\alpha}\right],$$

where the expectations are taken with respect to the same random factors as in Assumption 2.

Notably, again the independence of the false discoveries is not required here.

*Proof.* It follows from Assumption 3 that for $j \leq l \in \{1, \ldots, k+1\}$

$$P\left(R_\delta = j \,\middle|\, \delta \in \bar{\mathcal{O}} \cap \bar{\Phi}\,,\, \#\left(\Pi_\delta \cap \Delta\right) = l\,,\, K_\alpha\right) = P\left(R_\delta = k+1 \,\middle|\, \delta \in \bar{\mathcal{O}} \cap \bar{\Phi}\,,\, \#\left(\Pi_\delta \cap \Delta\right) = l\,,\, K_\alpha\right).$$

Multiplying both sides by $P\left(\delta \in \bar{\mathcal{O}} \cap \bar{\Phi}\,,\, \#\left(\Pi_\delta \cap \Delta\right) = l \,|\, K_\alpha\right)$ we get

$$P\left(\delta \in \bar{\mathcal{O}} \cap \bar{\Phi}\,,\, \#\left(\Pi_\delta \cap \Delta\right) = l\,,\, R_\delta = j \,|\, K_\alpha\right) = P\left(\delta \in \bar{\mathcal{O}} \cap \bar{\Phi}\,,\, \#\left(\Pi_\delta \cap \Delta\right) = l\,,\, R_\delta = k+1 \,|\, K_\alpha\right).$$

Therefore for $l \in \{1, \ldots, k+1\}$

$$\begin{aligned}
l \cdot E(N_l \,|\, K_\alpha) &= \sum_{\delta \in \bar{\mathcal{O}}} l \cdot P\left(\#\left(\Pi_\delta \cap \Delta\right) = l\,,\, R_\delta = k+1 \,|\, K_\alpha\right) \\
&\geq \sum_{\delta \in \bar{\mathcal{O}}} l \cdot P\left(\delta \in \bar{\mathcal{O}} \cap \bar{\Phi}\,,\, \#\left(\Pi_\delta \cap \Delta\right) = l\,,\, R_\delta = k+1 \,|\, K_\alpha\right) \\
&= \sum_{\delta \in \bar{\mathcal{O}}} \sum_{j=1}^{l} P\left(\delta \in \bar{\mathcal{O}} \cap \bar{\Phi}\,,\, \#\left(\Pi_\delta \cap \Delta\right) = l\,,\, R_\delta = j \,|\, K_\alpha\right) \\
&= \sum_{\delta \in \bar{\mathcal{O}}} P\left(\delta \in \bar{\mathcal{O}} \cap \bar{\Phi}\,,\, \#\left(\Pi_\delta \cap \Delta\right) = l\,,\, \delta \in \mathcal{O} \,|\, K_\alpha\right) \\
&= \sum_{\delta \in \bar{\mathcal{O}}} E\left(1_{\{\delta \in \mathcal{O} \cap \Phi\,,\, \#(\Pi_\delta \cap \Delta) = l\}} \,|\, K_\alpha\right) \\
&= E(\mathring{N}_l \,|\, K_\alpha).
\end{aligned}$$

Therefore,

$$\begin{aligned}
E(\widehat{\mathrm{FDP}}^{*k}_{\mathcal{E}\,\cup\,\mathcal{O}} \,|\, K_\alpha) &= E\left[\frac{N_\mathcal{E} + \sum_{l=1}^{k+1} l \cdot N_l}{K_\alpha} \,|\, K_\alpha\right] \\
&= \frac{E(N_\mathcal{E} \,|\, K_\alpha) + \sum_{l=1}^{k+1} l \cdot E(N_l \,|\, K_\alpha)}{K_\alpha} \\
&\geq \frac{E(N_\mathcal{E} \,|\, K_\alpha) + \sum_{l=1}^{k+1} E(\mathring{N}_l \,|\, K_\alpha)}{K_\alpha} \\
&= \frac{E(N_\mathcal{E} + \mathring{N}_\mathcal{O} \,|\, K_\alpha)}{K_\alpha} = E\left[\frac{N_\mathcal{E} + \mathring{N}_\mathcal{O}}{K_\alpha} \,|\, K_\alpha\right].
\end{aligned}$$

Taking expectation on both sides completes the proof. □

Like the combined estimate, the matched method provides an average upper bound on the FDP (and hence, when averaged, on the FDR). The advantage of the matched method is that it typically provides a smaller, i.e., tighter upper bound. To see why this is the case, consider the paired estimation specialized to the peptide-level analysis as in Equation (4), which coincides with Equation (S3) with $k = 1$, $N_1 = N_{\mathcal{E} \geq s > \mathcal{T}}$ and $N_2 = N_{\mathcal{E} > \mathcal{T} \geq s}$.

We can decompose $N_{\mathcal{E}>\mathcal{T}\geq s}$ (and the analogously defined $N_{\mathcal{T}>\mathcal{E}\geq s}$) into the sum of those cases where the paired target is in the sample (true discovery) $\check{N}_{\mathcal{E}>\mathcal{T}\geq s}$, and the cases where it is missing (false discovery) $\mathring{N}_{\mathcal{E}>\mathcal{T}\geq s}$:

$$
\begin{aligned}
N_{\mathcal{E}>\mathcal{T}\geq s} &= \check{N}_{\mathcal{E}>\mathcal{T}\geq s} + \mathring{N}_{\mathcal{E}>\mathcal{T}\geq s} \\
N_{\mathcal{T}>\mathcal{E}\geq s} &= \check{N}_{\mathcal{T}>\mathcal{E}\geq s} + \mathring{N}_{\mathcal{T}>\mathcal{E}\geq s}.
\end{aligned}
$$

It follows from Assumption 3 that $E(\mathring{N}_{\mathcal{E}>\mathcal{T}\geq s} \,|\, K_\alpha) = E(\mathring{N}_{\mathcal{T}>\mathcal{E}\geq s} \,|\, K_\alpha)$, but clearly for any reasonable score function, $E(\check{N}_{\mathcal{E}>\mathcal{T}\geq s} \,|\, K_\alpha) < E(\check{N}_{\mathcal{T}>\mathcal{E}\geq s} \,|\, K_\alpha)$: for peptides that are present in the sample we expect the target to score higher than its paired entrapment. Putting this together we have $E(N_{\mathcal{E}>\mathcal{T}\geq s} \,|\, K_\alpha) < E(N_{\mathcal{T}>\mathcal{E}\geq s} \,|\, K_\alpha)$ and therefore

$$
\begin{aligned}
E\left(\frac{N_{\mathcal{E}} + N_{\mathcal{E}\geq s>\mathcal{T}} + 2 \cdot N_{\mathcal{E}>\mathcal{T}\geq s}}{K_\alpha} \,\Big|\, K_\alpha\right) &< E\left(\frac{N_{\mathcal{E}} + N_{\mathcal{E}\geq s>\mathcal{T}} + N_{\mathcal{E}>\mathcal{T}\geq s} + N_{\mathcal{T}>\mathcal{E}\geq s}}{K_\alpha} \,\Big|\, K_\alpha\right) \\
&= E\left(\frac{N_{\mathcal{E}} + N_{\mathcal{E}}}{K_\alpha} \,\Big|\, K_\alpha\right),
\end{aligned}
$$

demonstrating that, on average, the paired estimate is smaller than the combined one.

### S3 The sample entrapment method can underestimate or overestimate the FDP

The sample entrapment estimation method (3) aims to estimate the FDP only among the original target discoveries rather than among the combined list of target+entrapment discoveries. Aside from the fact that it does not fall under our above formal definition, this approach is inherently problematic because the analyzing tool is asked to control the FDR in the target+entrapment list of discoveries while the evaluation is done only on the original target discoveries. As a result, while as explained next, the sample entrapment method typically underestimates the FDP, it can also potentially overestimate the FDP. Therefore, this method cannot be used to show that a tool controls the FDR, nor that it does not control the FDR. The related fallacy of trying to control the FDR in a subset by controlling it in the larger set has already been pointed out in the literature [2, 5].

Indeed, typically the FDP among the discoveries from the target $\mathcal{T}$ will be significantly lower than among the target+entrapment list of discoveries, and hence (3) will typically underestimate the FDP among the latter discoveries. Critically, it is the latter FDP that the search tool is actually trying to control, so typically the sample estimation will underestimate the true FDP.

To demonstrate that (3) can in some extreme cases overestimate the FDP, we find it simpler to assume that the analyzing tool aims to report peptides in DDA data. Consider a hypothetical case, where every original target peptide in $\mathcal{T}$ produces a very good spectrum that creates a perfect PSM. Suppose that, in addition to those perfect spectra, the dataset also contains a large number of other spectra that are foreign to $\mathcal{T}$. Because the native spectra will mostly be optimally matched to the original target peptides they were generated from, the entrapment peptides will overwhelmingly be optimally matched to the foreign spectra. Inevitably, some of those entrapment peptides will score fairly high and make it to the target+entrapment discovery list. Consequently, we will end up with $N_{\mathcal{E}} > 0$ and assuming $r = 1$ the sample method in Equation (3) will yield

$$\widehat{\mathrm{FDP}}_{\mathcal{T}} = \frac{N_{\mathcal{E}}}{N_{\mathcal{T}}} > \frac{N_{\mathcal{E}}}{N_{\mathcal{T}} + N_{\mathcal{E}}}.$$

Notably, the expression on the right is the real FDP in the combined list of discoveries because there are no false matches among the original target, so the sample method overestimates the true FDP in this case.

Other versions of (3) that aim to estimate the FDP among the discoveries from the original target $\mathcal{T}$ have been used as well. For example, Yu *et al.* [7] estimate this FDP by $\frac{N_{\mathcal{E}}}{N_{\mathcal{T}}} \frac{M_{\mathcal{T}} - N_{\mathcal{T}}}{M_{\mathcal{E}}}$, where $M_{\mathcal{T}}$ and $M_{\mathcal{E}}$ are the number of peptides in the original target and entrapment databases. Much of the above criticism applies to this estimator as well: it can overestimate as well as underestimate the FDP. For example, if the tool reports that it detected all the original target peptides, then this estimate is 0, but of course, this will typically underestimate the FDP. Similarly, if all the original target peptides are present in the sample then the actual FDP is 0, but this estimate will typically be positive.

### S4 Potential pitfalls of using foreign entrapment sequences

Critical to all methods of FDP estimation is the ratio $r$ of entrapment-to-original-target sequences. When using shuffled entrapment every original target sequence creates exactly $r$ shuffled entrapment sequences, each with the same composition as the target's, and therefore there is no ambiguity about that ratio. In contrast, consider using a foreign entrapment in peptide-level analysis. The natural choice for $r$ is the ratio of the number of entrapment-to-original-target peptides. However, it is conceivable that the entrapment peptides would have very different amino acid composition and length distributions compared with those of the original target peptides. Such differences can cast doubts over the adequacy of that simple ratio.

#### S4.1 Selecting the "right" evolutionary distance

An issue related to the last point is the critical selection of the foreign species: choose species that are at a very large evolutionary distance to your target species and you risk creating entrapment sequences that do not offer a realistic competition to the incorrect target discoveries they need to model. In this case you are likely to underestimate the true FDP when using the combined estimation. Conversely, if your entrapment includes a species that is a close relative of your target, then you run the risk of overestimating the FDP, particularly when considering sample peptides whose exact matches are missing from the original target database, say due to unexpected modifications or genetic variations.

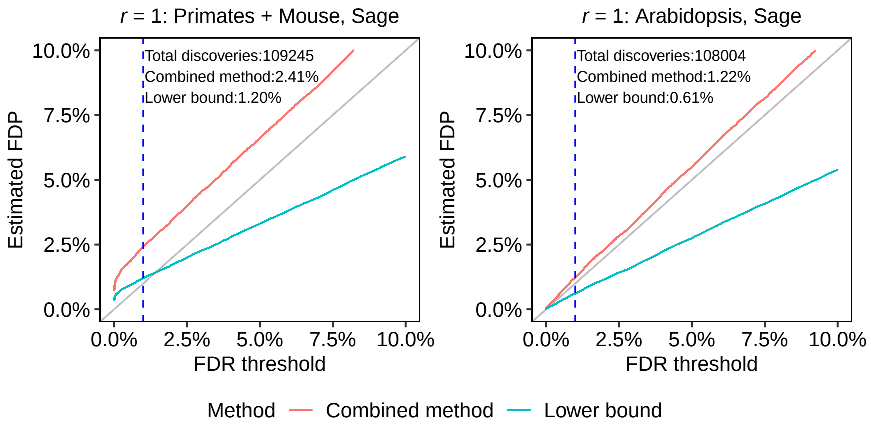


Figure S1: **Varying the evolutionary distance of the peptide-level foreign entrapment with Sage applied to the HEK293 dataset ($r = 1$).** The dashed vertical lines are at the 1% FDR threshold, as are the numbers reported in text in the panels.

We next demonstrate this point through a few sets of entrapment experiments we conducted, where in each set we use the same analysis tool and the same dataset, while varying the evolutionary distance of the foreign entrapment sequences even as we keep a fixed $r = 1$.

Figure S1 presents the different conclusions we can draw when evaluating Sage's peptide-level analysis of the human-sampled HEK293 DDA dataset using two different foreign entrapment resources (both described in Section 4.5). As expected, when using a mixture of (non-ape) primates and mouse proteins for our foreign entrapment (left panel), Sage's estimated FDPs are substantially higher than when using Arabidopsis for the entrapment (right panel). Because we cannot use the paired estimation when using a foreign entrapment, the combined estimation is the only way to establish FDR control in this case. However, the combined curve is noticeably above the diagonal in the left panel. In particular, it estimates the FDP for the 1% FDR threshold at a rather high 2.4%, thus calling into question Sage's FDR control had we only used that entrapment experiment. In contrast, using the Arabidopsis peptides the combined method is only marginally above the diagonal (right panel). Recalling the general conservative nature of the combined method this raises the possibility that using now the much more evolutionary distant Arabidopsis we might be slightly underestimating the FDP in this case.

We observe the same trends when using foreign entrapments to evaluate DIA-NN's analysis of the human-sampled PXD042704 (human-astral) DIA dataset. Specifically, Supplementary Figure S2's comparison of using the Primates+Mouse vs. Arabidopsis entrapment sequences to evaluate DIA-NN's precursor-level analysis looks qualitatively very similar to the corresponding peptide-level analysis of Sage in Figure S1, so the same comments apply here.

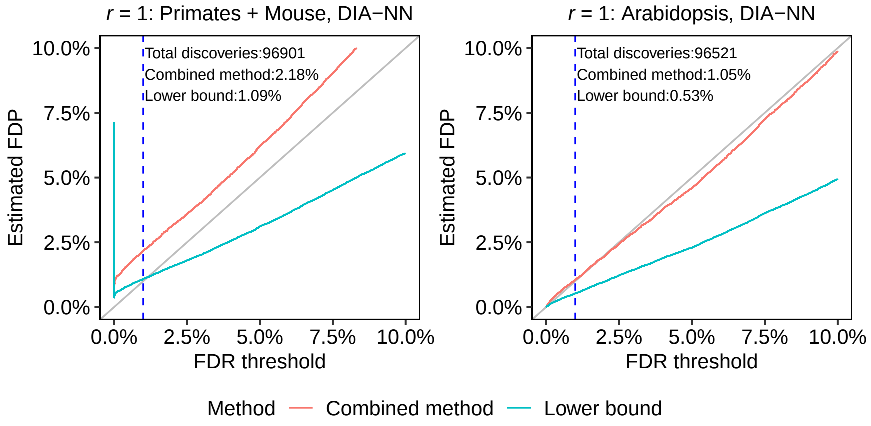


Figure S2: **Varying the evolutionary distance of the precursor-level foreign entrapment with DIA-NN applied to the DIA dataset PXD042704 (human-astral) with $r = 1$.** The dashed vertical lines are at the 1% FDR threshold, as are the numbers reported in text in the panels.

Similarly, the protein-level entrapment analyses depicted in Figure S3 again show that all estimated FDPs increase as the evolutionary distance to the human sample decreases.

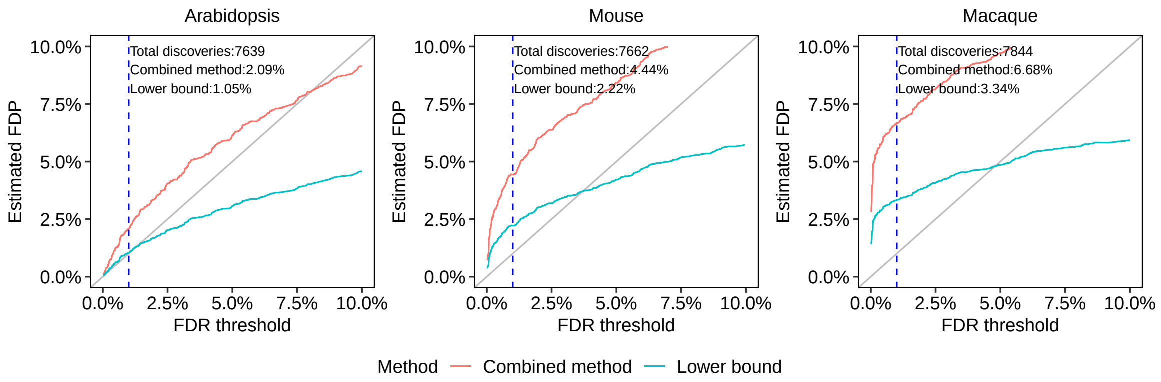


Figure S3: **Varying the evolutionary distance of the protein-level foreign entrapment with DIA-NN applied to the DIA dataset PXD042704 (human-astral) with $r = 1$.** The dashed vertical lines are at the 1% FDR threshold, as are the numbers reported in text in the panels.

#### S4.2 Foreign entrapments can be compromised by contamination

Finally, it is important to note that an entrapment experiment that relies on foreign sequences can be led astray by contamination. Indeed, compare the results of the two entrapment experiments summarized in Figure S4, where in both cases Sage peptide-level analysis was applied to the HEK293 dataset using two different foreign entrapments, both constructed so that $r = 1$: one made of yeast + Arabidopsis peptides (left panel) and the other made of Arabidopsis-only peptides (right panel). Clearly, adding the yeast peptides to the entrapment database significantly increases both the combined and the lower bound estimates of the FDP.

Looking more closely, we find that at the 1% FDR threshold Sage reported 108,867 peptides, of which 1,409 were entrapment peptides corresponding to a combined estimate of 2.59% ($r = 1$), and a lower bound of 1.29% on the FDP. Notably, 897 of the 1,409 discovered entrapment peptides (63.7%) were yeast peptides even though only 289,258 of the 1,389,436 entrapment peptides (20.8%) came from yeast. Statistically this is a highly significant enrichment of the yeast peptides among the discovered entrapment peptides (two-sided Fisher Exact p-value $< 2.2e\text{-}16$). Unlike the examples in the previous section, yeast of course is not significantly closer to human than Arabidopsis, thus yeast contamination remains the most likely explanation.

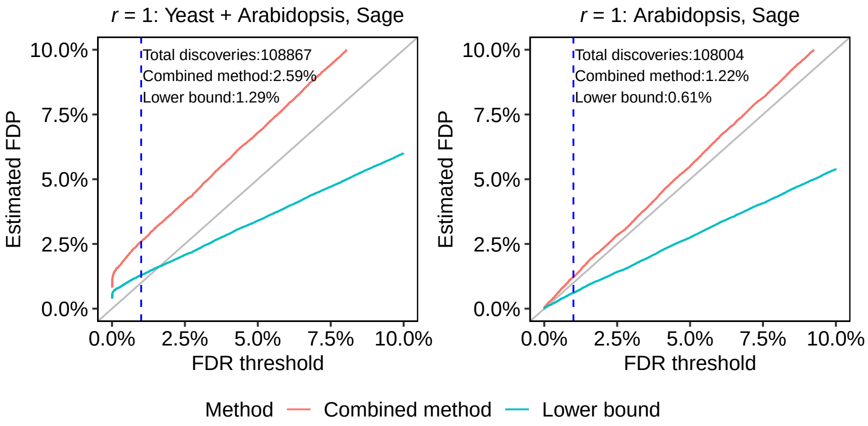


Figure S4: **Foreign entrapments can be biased by contamination.**

Consistently, searching the HEK293 MS/MS dataset against a protein database containing human and yeast proteins using Sage, we found that 1,510 out of the 111,376 peptides reported at 1% FDR match only to yeast proteins (a further 486 peptides match both the yeast and human proteins). Factoring in the ratio of human to yeast peptides this 1.4% of yeast peptides is much larger than expected. Moreover, if we specifically focus on the run with ID 02B, we find that 961 of the 9,390 peptides reported at 1% FDR match only to yeast proteins (a further 103 peptides match both the yeast and human proteins). This 10.2% yeast content is a fairly strong indication of some yeast contamination in that specific run.

### S5 A survey of notable entrapment experiments

We conducted an extensive literature review looking at some prominent publications that used entrapment experiments. The results are summarized in Table 1, and here we discuss a few of those publications in more detail.

In [3] the authors estimated the FDP as $\frac{N_{\mathcal{E}} \cdot 1/r}{N_{\mathcal{T}} + N_{\mathcal{E}} \cdot 1/r}$, which does not seem to be well motivated. Applying this formula to the numbers they reported using—$N_{\mathcal{T}} = 8,867$, $N_{\mathcal{E}} = 297$ (*E. coli* peptides) and $r = 48,131/29,333$–we obtain the 0.02 estimated FDP that they reported. Regardless of the fact that this estimate is already higher than the 1% FDR threshold, if you use the combined estimation method, Equation (1), then you get an estimate higher than 0.05, or 5 times higher than the threshold. Moreover, even the lower bound, Equation (2), is larger than 0.03, or 3 times higher than the threshold, casting doubt over the claim that Specter controls the FDR.

In [4] the authors indicated that they used $r = 1.176$ and claim that the estimated FDPs (referred to as "external FDR") were in very good agreement with the corresponding thresholds ("internal FDR"). However, they used the sample method, which as we argued is invalid. Had the authors used the valid combined method, then taking $r$ into account, the 1% FDP estimate they computed would have translated to 2.2%. Because this is not a lower bound it does not demonstrate MaxDIA fails to control the FDR, but at the same time this experiment does not provide evidence to support the fact that it controls the FDR.

In [7] the authors also used the invalid sample method to estimate the FDP among the quantified proteins reported by three different DIA tools. Using their reported values, $r = (6060 + 4401 + 16224)/20407 \approx 1.31$ the estimated FDPs (at 1% FDR) they reported for DIA-NN library-free, FP-MSF, and FP-MSF hybrid pipelines of 1.2%, 1.9%, and 1.8% respectively, correspond to 2.7%, 4.3% and 4.1% respectively if we use the valid combined method. Again, this analysis does not prove that these three tools do not control the FDR, but this experiment cannot be taken as evidence of proper FDR control.

In [1] the lower bound was correctly used to argue questionable FDR control in the case of MaxDIA, but it was also incorrectly used to argue proper FDR control of Spectronaut and DIA-NN. As we mentioned, the lower bound cannot be used as evidence of proper FDR control. For example, the authors report that using an *in silico* entrapment setup, DIA-NN's estimated FDP at 1% FDR was 1.77%, which they found acceptable. However, keeping in mind that this is a lower bound and that given that the entrapment part consisted of 16,202 *Arabidopsis* protein sequences and the original target was made of 17,082 mouse and 6,730 yeast protein sequences, we have $r \approx 0.68$. Therefore, using the same entrapment setup, the combined method estimates the FDP at 2.6%. The real FDP is somewhere in between, but it is highly questionable whether these results can be taken as evidence for proper FDR control.

The paper describing the recently released AlphaPept refers to some entrapment experiments, but the method is not fully described [6]. Looking at the accompanying code they released, we found that they were estimating the FDP as $N_{\mathcal{E}}/(N_{\mathcal{E}} + N_{\mathcal{E}} + N_{\mathcal{D}})$, where $N_{\mathcal{D}}$ is the number of decoys above the cutoff. This estimate is even smaller than the lower bound on the FDP (Equation (2)); hence, it cannot possibly be used as evidence of valid FDR control.

### S6 Supplementary Algorithms

**Algorithm 1** Entrapment database generation for precursor/peptide-level analysis using random shuffling of target peptides. Note that the algorithm can be readily modified to keep track of the pairing/matching information.

**Input:**
    $P_{\text{target}}$ - Original target proteins
    $min\_len, max\_len$ - Minimum and maximum peptide lengths (7 and 35 amino acids)
    $r$ - Number of distinct random entrapment peptides to generate
    $max\_attempts = 20 + r$ - Maximum attempts to generate distinct random peptides
**Output:**
    $P_{\text{database}}$ - Processed entrapment peptide database: original target peptides and entrapment peptides

**1. Peptide digestion:**
**for** each protein $p \in P_{\text{target}}$ **do**
    $P_{\text{target\_peptides}} \leftarrow \text{in\_silico\_digest}(p, \text{trypsin}, \text{missed\_cleavage} = 1)$
**end for**

**2. Filter peptides by length:**
$P_{\text{target\_peptides}} \leftarrow \{\text{peptide} \in P_{\text{target\_peptides}} \mid 7 \leq \text{len(peptide)} \leq 35\}$

**3. Generate random entrapment peptides:**
**for** each peptide $p_{\text{target}} \in P_{\text{target\_peptides}}$ **do**
    $P_{\text{shuffled}} \leftarrow \emptyset$
    attempts $\leftarrow 0$
    **while** $|P_{\text{shuffled}}| < r$ **and** attempts < max_attempts **do**
        Shuffle $p_{\text{target}}$, keeping the C-terminal fixed
        **if** the shuffled peptide is distinct from all peptides in $P_{\text{target\_peptides}}$ and $P_{\text{shuffled}}$ **then**
            Add shuffled peptide to $P_{\text{shuffled}}$
        **else**
            Retry shuffling
        **end if**
        Increment attempts
    **end while**
    **if** $|P_{\text{shuffled}}| < r$ **then**
        Remove $p_{\text{target}}$ from $P_{\text{target\_peptides}}$
    **end if**
**end for**

**4. Update database:**
Add $P_{\text{shuffled}}$ to the database: $P_{\text{database}} \leftarrow P_{\text{target\_peptides}} \cup P_{\text{shuffled}}$

**5. Return the processed peptide database:**
**return** $P_{\text{database}}$

**Algorithm 2** Entrapment database generation for precursor/peptide-level analysis using peptides from foreign species

**Input:**
  $P_{\text{target}}$ - Original target proteins
  $P_{\text{foreign}}$ - Foreign species proteins
  $r$ - Desired ratio of entrapment-to-target peptides ($r \geq 1$)
  $min\_len, max\_len$ - Minimum and maximum peptide lengths (7 and 35 amino acids)
**Output:**
  $P_{\text{database}}$ - Peptide-level entrapment database

**1. Peptide digestion of target proteins:**
**for** each protein $p_{\text{target}} \in P_{\text{target}}$ **do**
  $P_{\text{target\_peptides}} \leftarrow \text{in\_silico\_digest}(p_{\text{target}}, \text{trypsin}, \text{missed\_cleavage} = 1)$
**end for**

**2. Peptide digestion of foreign proteins:**
**for** each protein $p_{\text{foreign}} \in P_{\text{foreign}}$ **do**
  $P_{\text{foreign\_peptides}} \leftarrow \text{in\_silico\_digest}(p_{\text{foreign}}, \text{trypsin}, \text{missed\_cleavage} = 1)$
**end for**

**3. Filter peptides by length:**
$P_{\text{target\_peptides}} \leftarrow \{\text{peptide} \in P_{\text{target\_peptides}} \mid 7 \leq \text{len(peptide)} \leq 35\}$
$P_{\text{foreign\_peptides}} \leftarrow \{\text{peptide} \in P_{\text{foreign\_peptides}} \mid 7 \leq \text{len(peptide)} \leq 35\}$

**4. Remove matching foreign peptides:**
**for** each peptide $p_{\text{foreign}} \in P_{\text{foreign\_peptides}}$ **do**
  **if** $p_{\text{foreign}} \in P_{\text{target\_peptides}}$ **then**
    Remove $p_{\text{foreign}}$
  **end if**
**end for**

**5. Calculate the number of foreign peptides needed:**
Let $n_{\text{target}} \leftarrow |P_{\text{target\_peptides}}|$
Let $n_{\text{foreign\_needed}} \leftarrow r \times n_{\text{target}}$

**6. Randomly select foreign peptides:**
**if** $n_{\text{foreign\_needed}} \leq |P_{\text{foreign\_peptides}}|$ **then**
  Randomly select $n_{\text{foreign\_needed}}$ peptides from $P_{\text{foreign\_peptides}}$
**else**
  Use all available foreign peptides
**end if**

**7. Update database:**
$P_{\text{database}} \leftarrow P_{\text{target\_peptides}} \cup P_{\text{foreign\_selected}}$

**8. Return the entrapment database:**
**return** $P_{\text{database}}$

#### S7 Additional Supplementary Figures

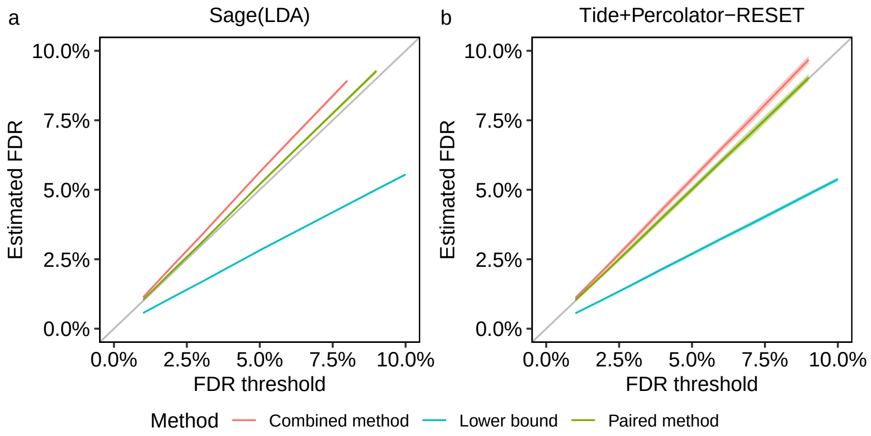


Figure S5: **Variability of the entrapment estimations of Tide's and Sage's on the HEK293 DDA dataset.** (a) Sage. (b) Tide. The entrapment-estimated FDR ($r = 1$) together with 95% coverage bands are plotted as a function of the given FDR threshold. The estimated FDR is the average of the estimated FDP over 10 sets of shuffled decoys and entrapment sets in Tide's case. Sage uses reversed peptides as decoys so we could only vary the entrapment sequences in that case, and hence its 95% coverage bands are visually indistinguishable from the estimates themselves. The FDP is estimated using three different entrapment methods.

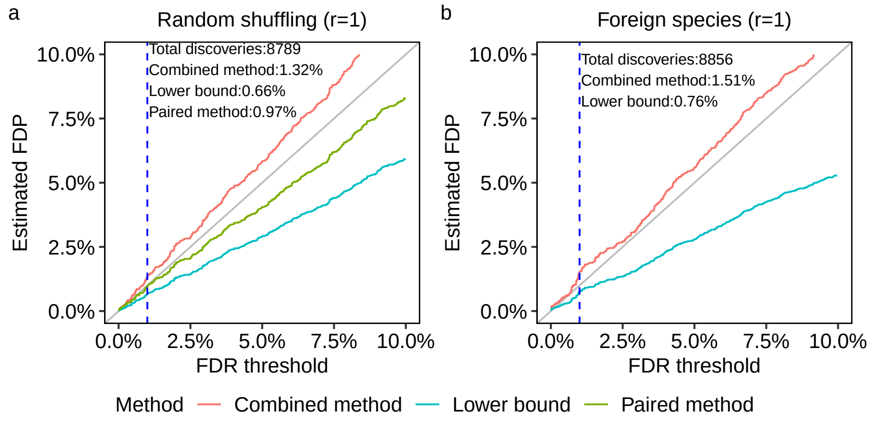


Figure S6: **Protein-level FDR control evaluation of MaxQuant on the HEK293 DDA dataset.** The protein-level analysis FDP was estimated using $r = 1$ and (a) shuffled entrapment sequences, as well as (b) foreign entrapment sequences (arabidopsis). Each panel plots the estimated FDP as a function of the FDR threshold. The dashed vertical lines are at the 1% FDR threshold, as are the numbers reported in text in the panels.

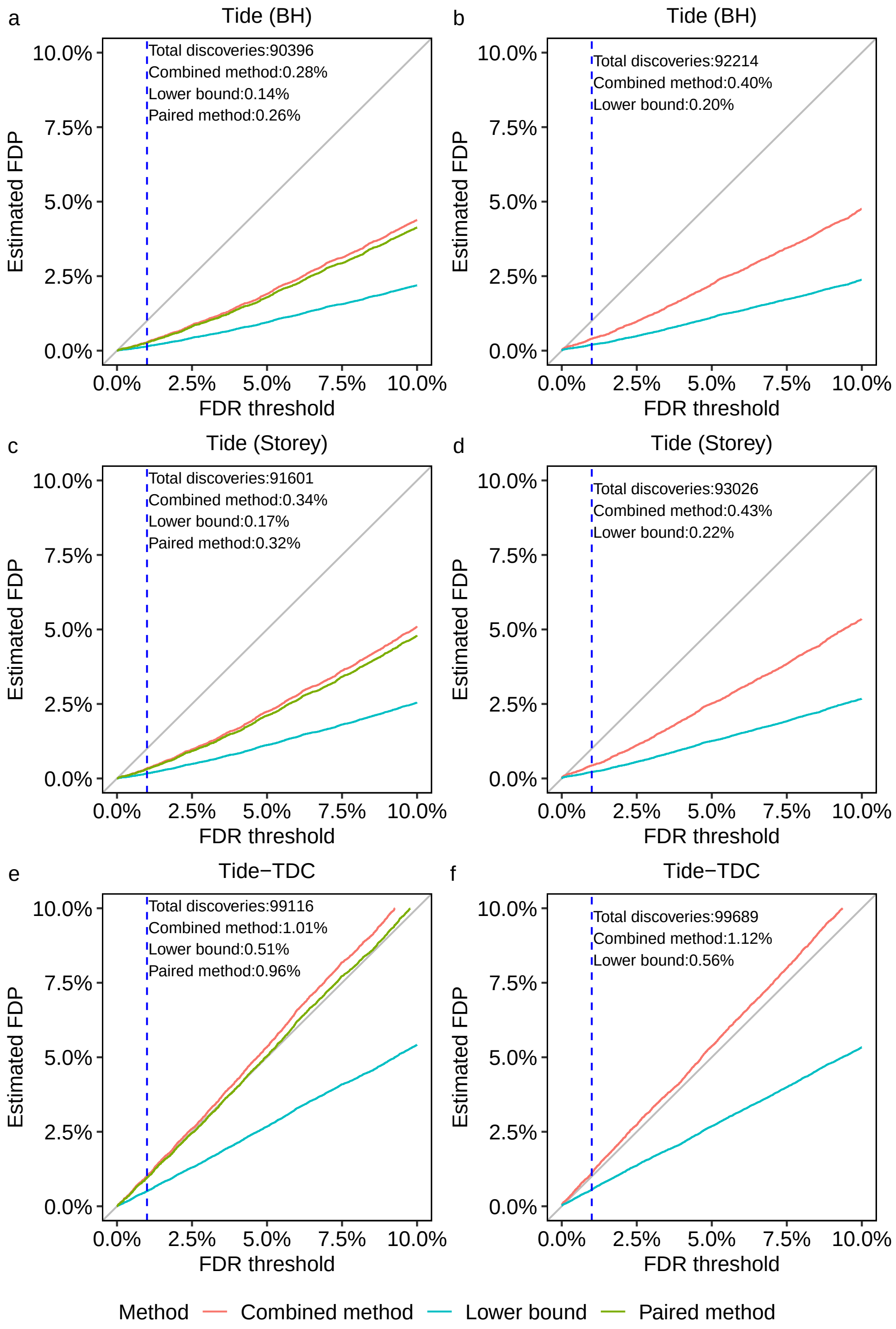


Figure S7: **Comparing multiple modes of controlling the FDR and entrapment-sequence generation.** The HEK293 dataset was analyzed by Tide with varied FDR control: empirically estimated p-values using BH (top row), using Storey (middle row), and using TDC in lieu of p-values (bottom row). The FDP was estimated using $r = 1$ with both shuffled entrapment sequences (left column) and foreign sequences (arabidopsis, right column). Each panel plots the estimated FDP as a function of the FDR threshold. The dashed vertical lines are at the 1% FDR threshold, as are the numbers reported in text in the panels.

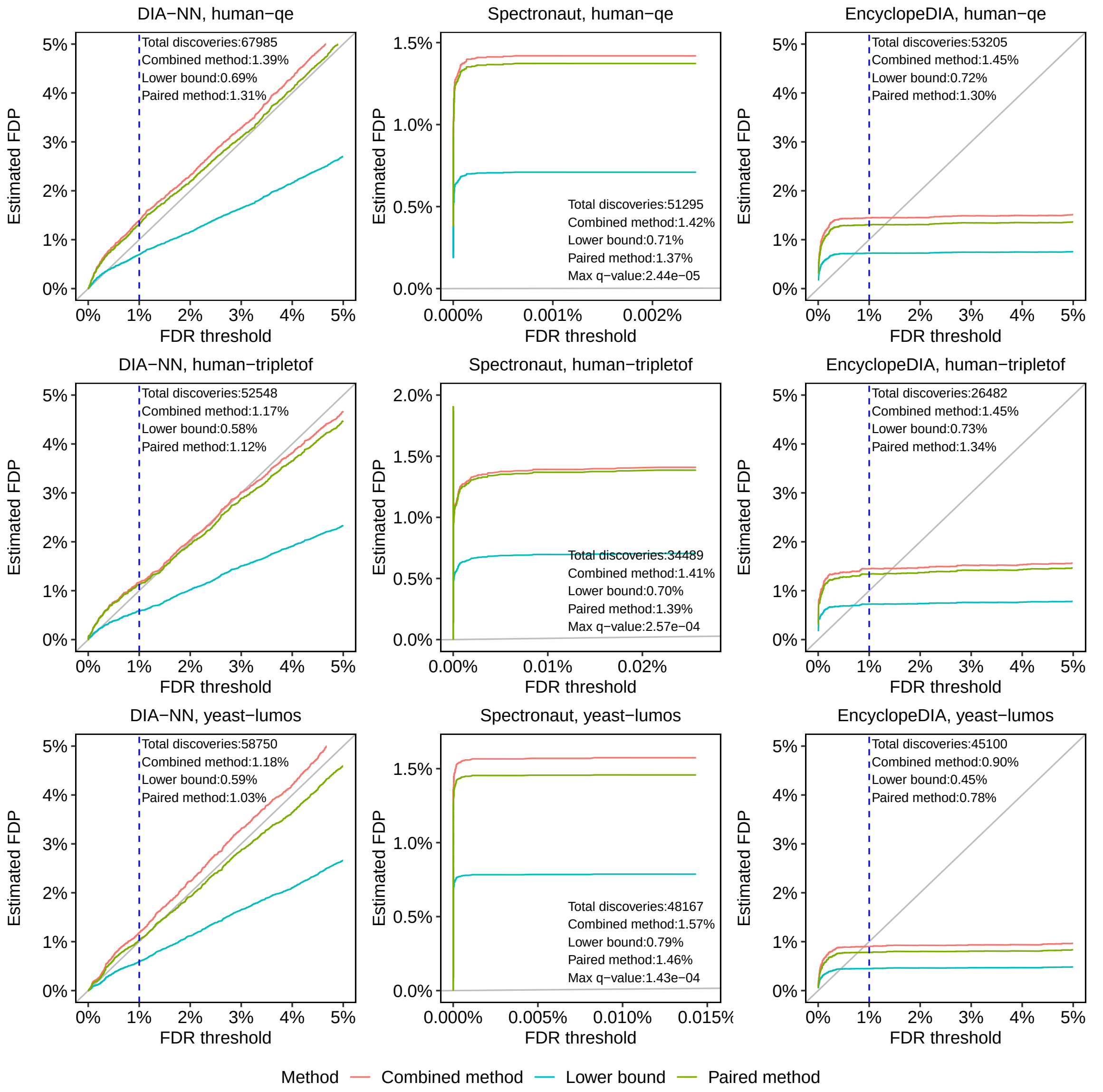


Figure S8: **Peptide or precursor-level FDR control evaluation of DIA search tools** In each panel, the empirical FDP was estimated for a given dataset (row) and search tool (column) using three different entrapment methods. Each panel plots the estimated FDP as a function of the FDR threshold. The dashed vertical lines are at the 1% FDR threshold, as are the numbers reported in text in the panels. The "Max q-value" value in each Spectronaut panel was the maximum reported q-value by the tool.

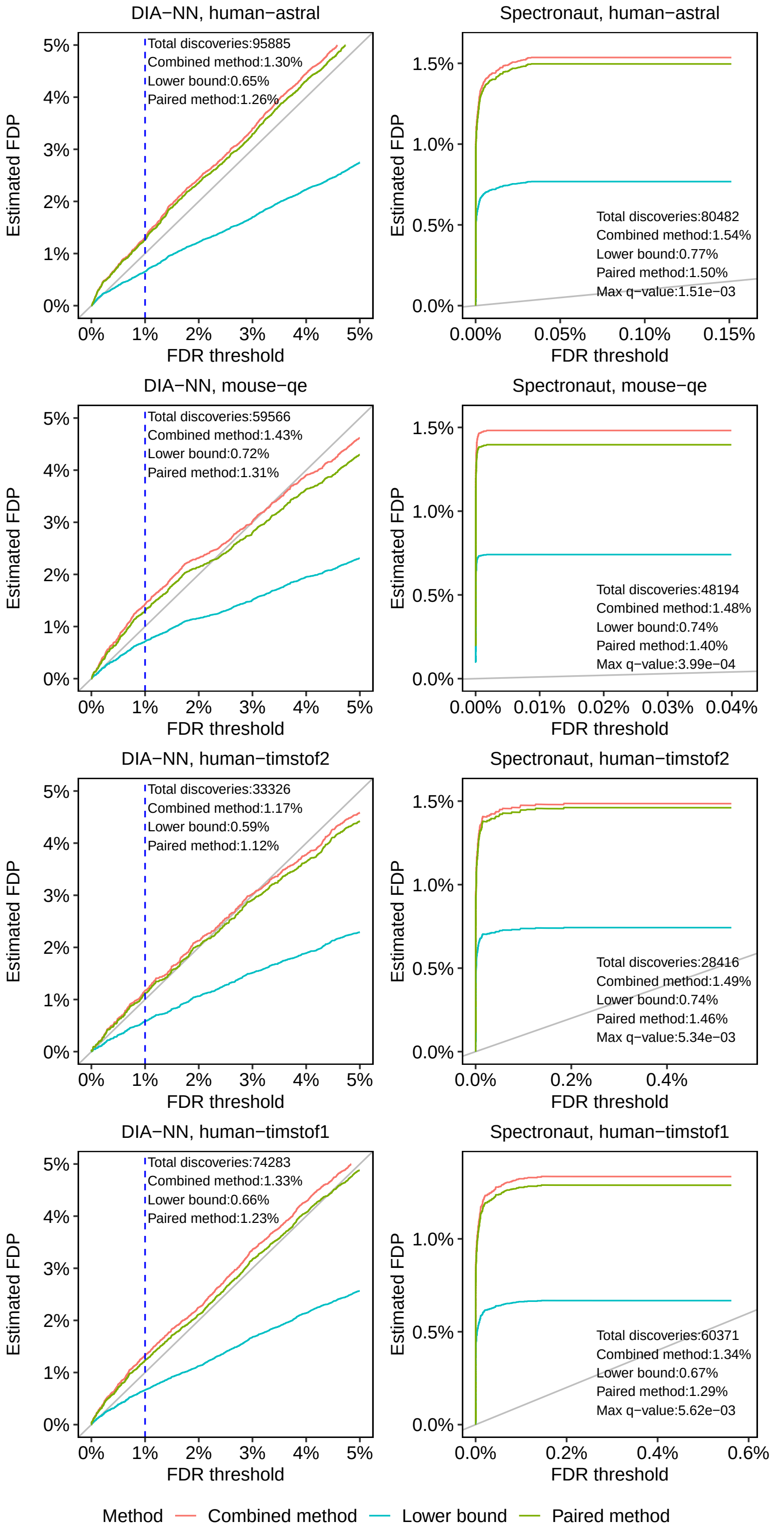


Figure S9: **Precursor-level FDR control evaluation of DIA search tools** In each panel, the empirical FDP was estimated for a given dataset (row) and search tool (column) using three different entrapment methods. Each panel plots the estimated FDP as a function of the FDR threshold. The dashed vertical lines are at the 1% FDR threshold, as are the numbers reported in text in the panels. The "Max q-value" value in each Spectronaut panel was the maximum reported q-value by the tool.

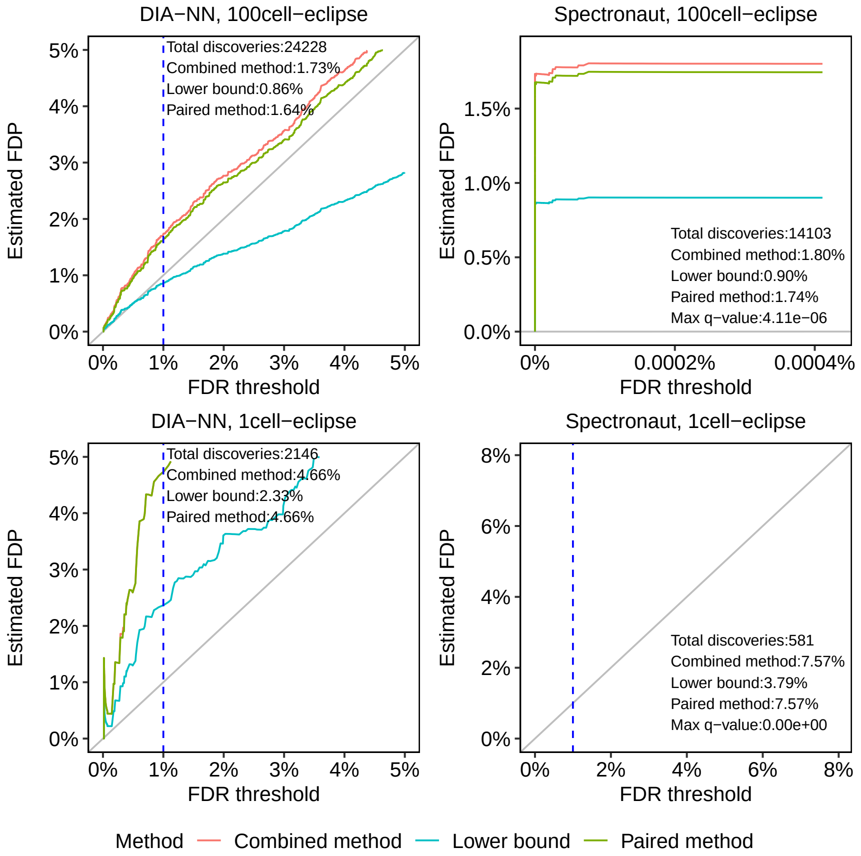


Figure S10: **Precursor-level FDR control evaluation of DIA search tools** In each panel, the empirical FDP was estimated for a given dataset (row) and search tool (column) using three different entrapment methods. Each panel plots the estimated FDP as a function of the FDR threshold. The dashed vertical lines are at the 1% FDR threshold, as are the numbers reported in text in the panels. The "Max q-value" value in each Spectronaut panel was the maximum reported q-value by the tool.

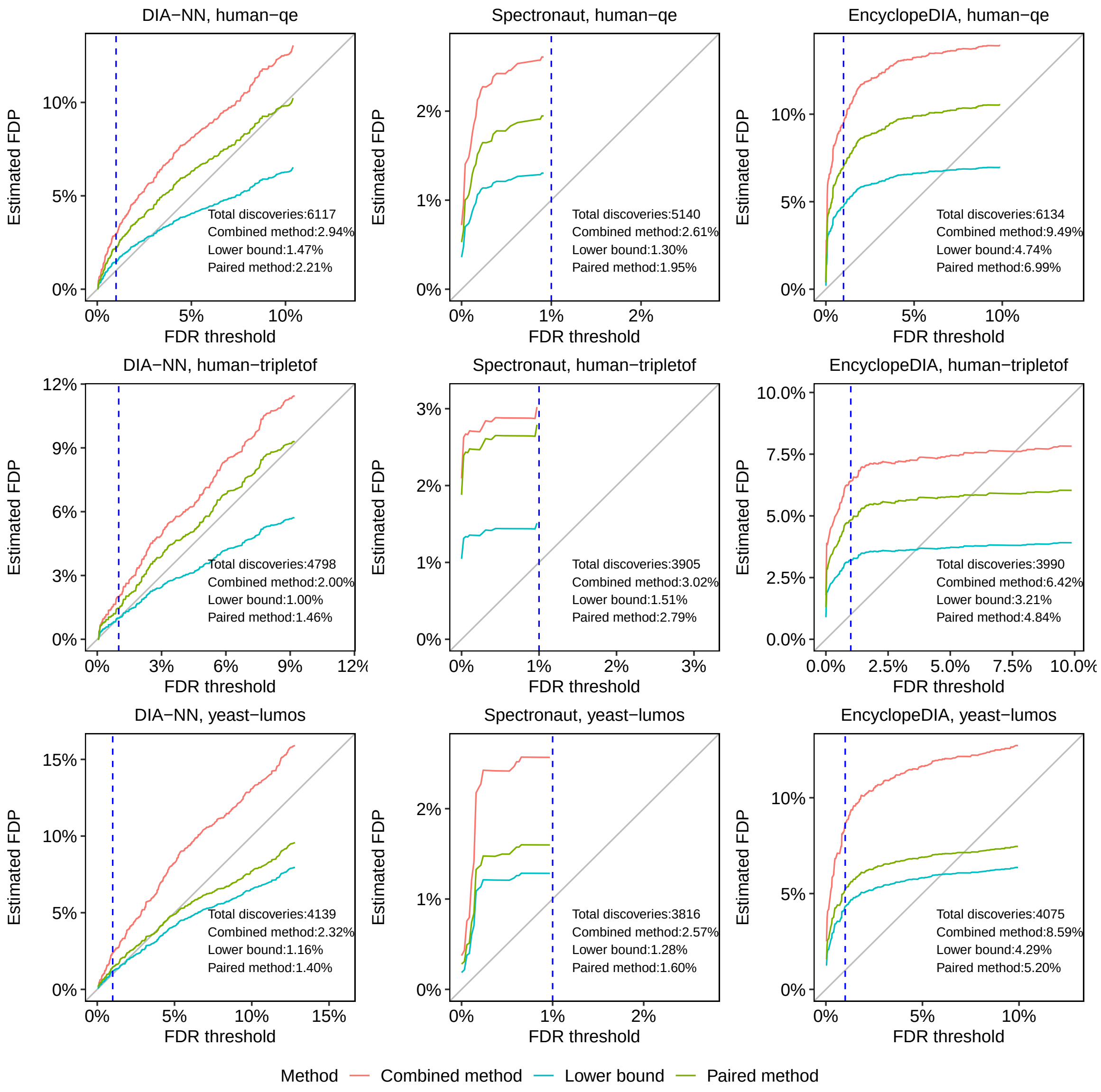


Figure S11: **Protein-level FDR control evaluation of DIA search tools** In each panel, the empirical FDP was estimated for a given dataset (row) and search tool (column) using three different entrapment methods. Each panel plots the estimated FDP as a function of the FDR threshold. The dashed vertical lines are at the 1% FDR threshold, as are the numbers reported in text in the panels.

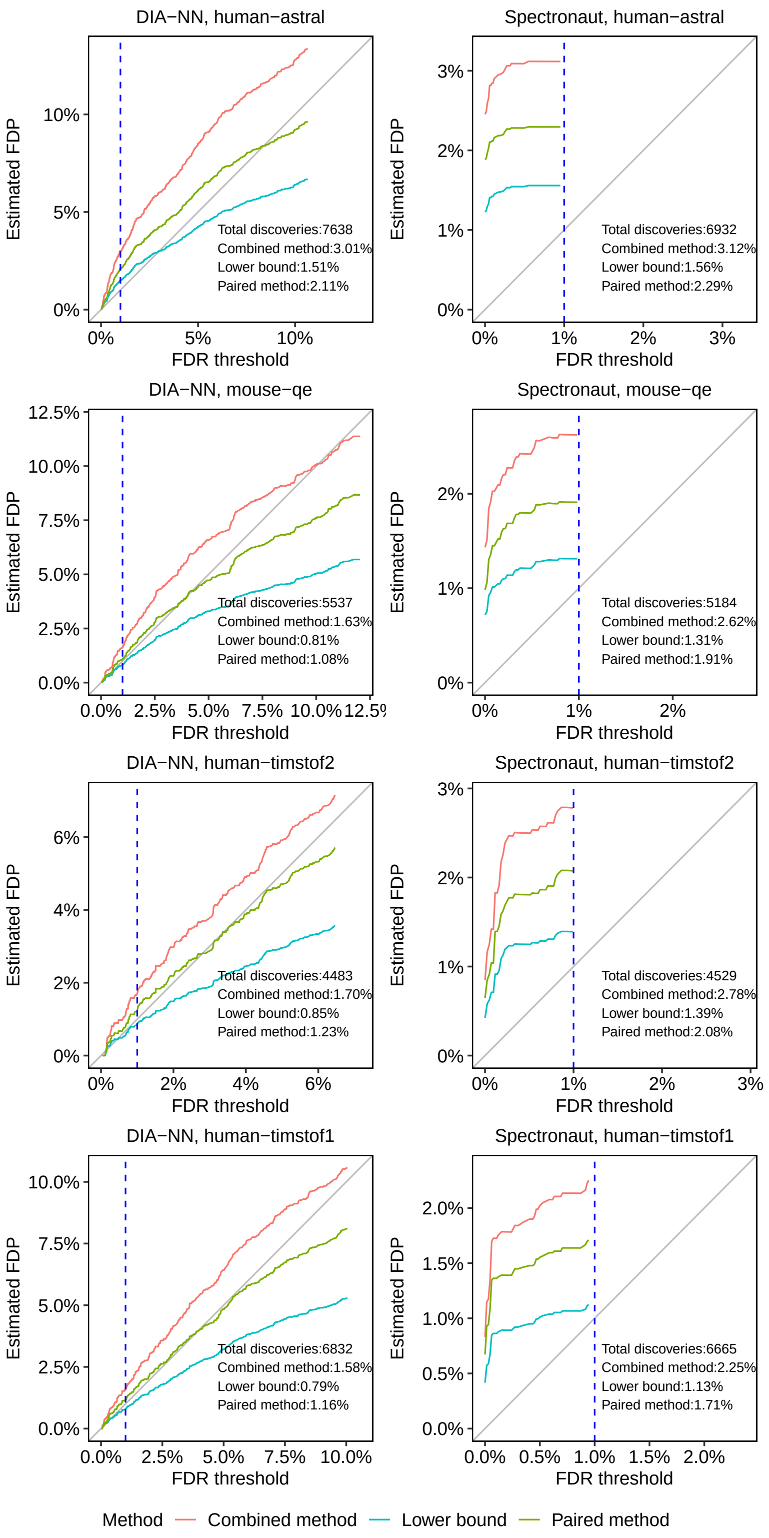


Figure S12: **Protein-level FDR control evaluation of DIA search tools** In each panel, the empirical FDP was estimated for a given dataset (row) and search tool (column) using three different entrapment methods. Each panel plots the estimated FDR as a function of the FDR threshold. The dashed vertical lines are at the 1% FDR threshold, as are the numbers reported in text in the panels.

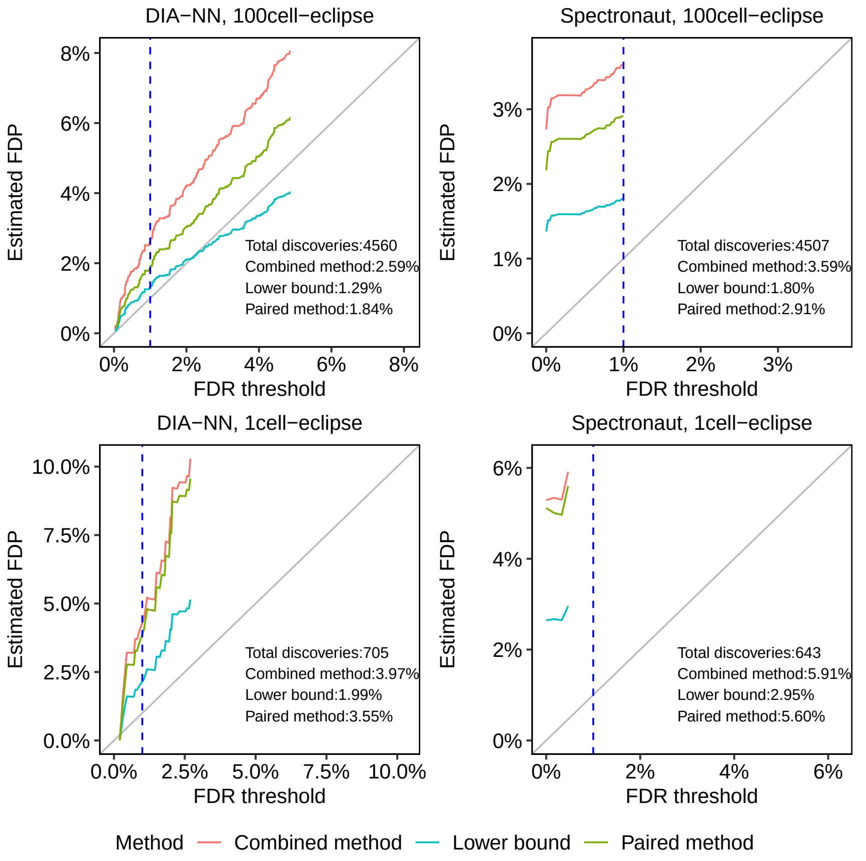


Figure S13: **Protein-level FDR control evaluation of DIA search tools** In each panel, the empirical FDP was estimated for a given dataset (row) and search tool (column) using three different entrapment methods. Each panel plots the estimated FDP as a function of the FDR threshold. The dashed vertical lines are at the 1% FDR threshold, as are the numbers reported in text in the panels.

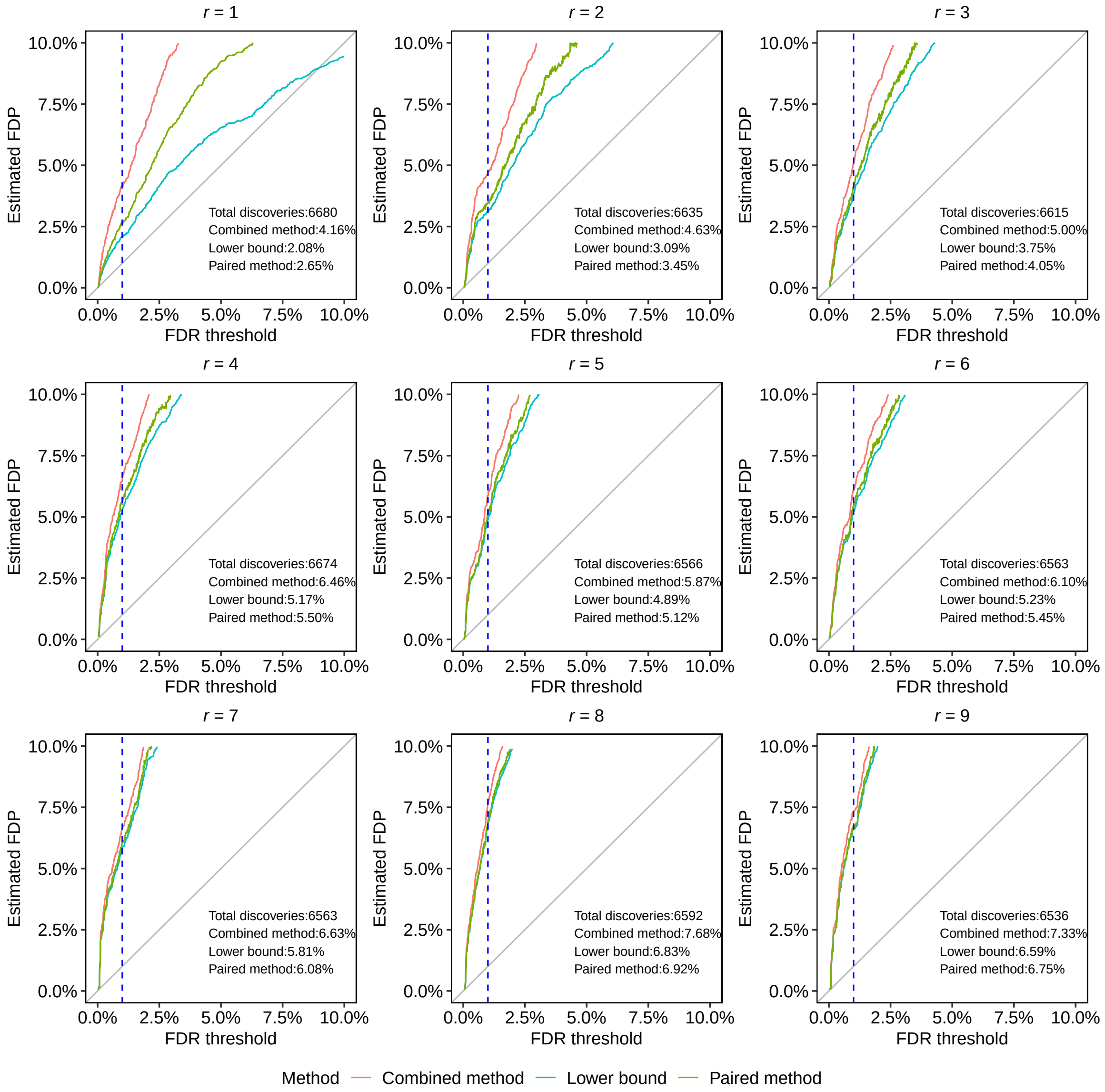


Figure S14: **DIA-NN struggles to control the FDR at the protein level as $r$ increases on the DIA dataset PXD034525 (human-lumos).**

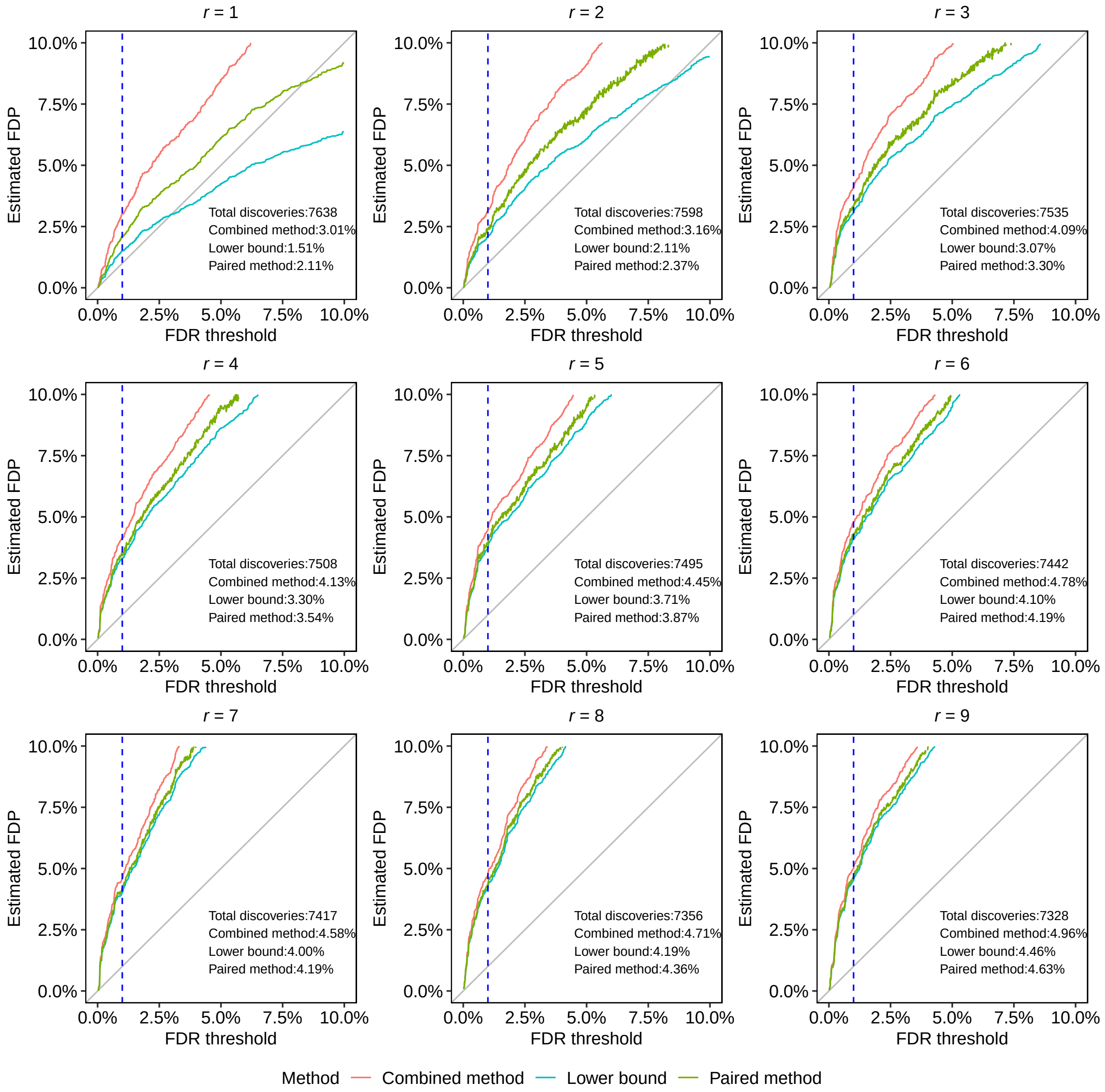


Figure S15: **DIA-NN struggles to control the FDR at the protein level as $r$ increases on the DIA dataset PXD042704 (human-astral).**

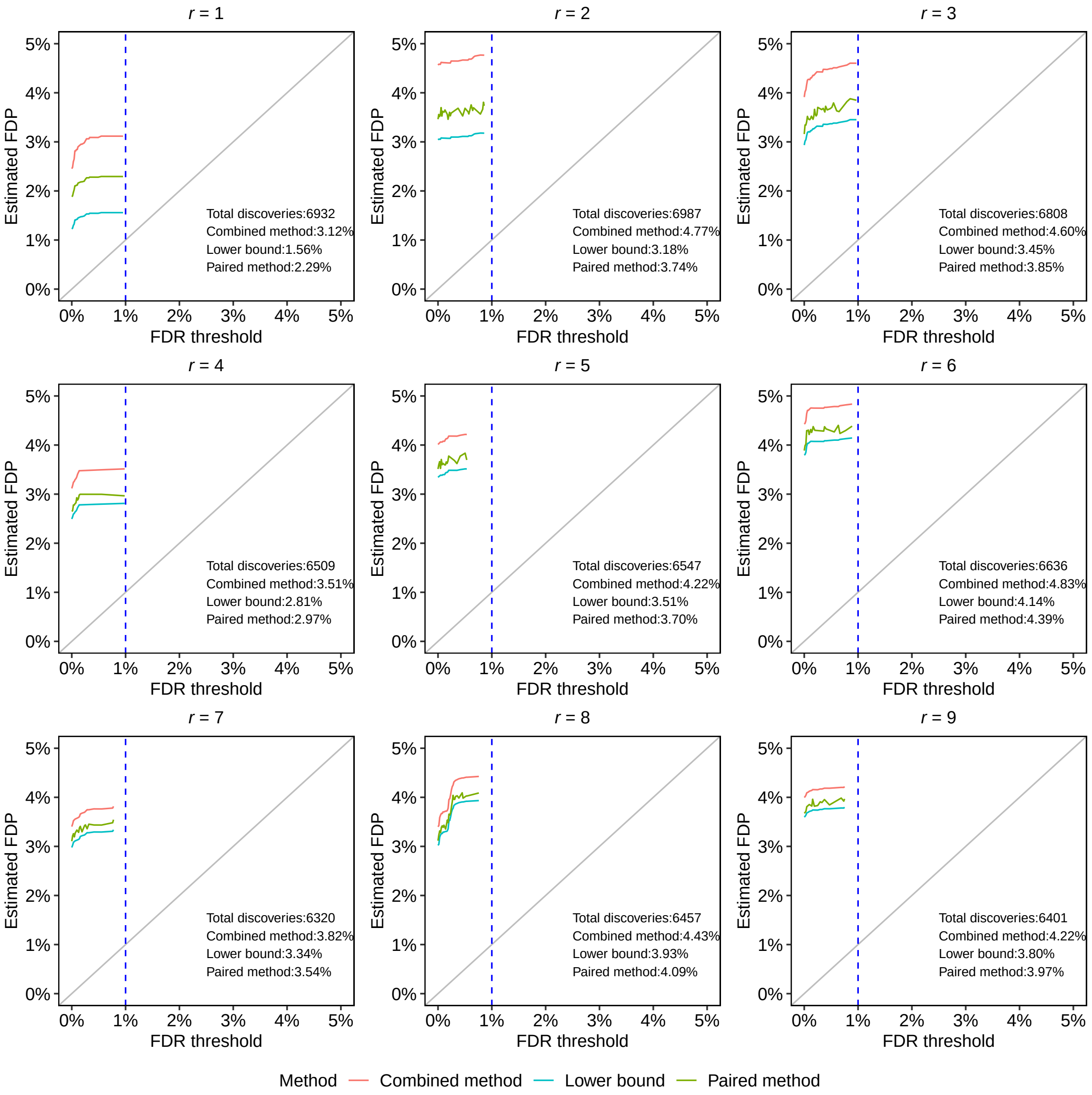


Figure S16: **Spectronaut struggles to control the FDR at the protein level as $r$ increases on the DIA dataset PXD042704 (human-astral).** As $r$ increases all methods report increasingly larger estimated FDP. Specifically, with $r = 6$ even the lower bound on the FDP among the 6,636 proteins discovered at 1% FDR is as high as 4.1%.

### References


[1] R. Lou, Y. Cao, S. Li, X. Lang, Y. Li, Y. Zhang, and W. Shui. Benchmarking commonly used software suites and analysis workflows for dia proteomics and phosphoproteomics. *Nature Communications*, 14(1):94, 2023.

[2] W. S. Noble and U. Keich. Response to "Mass spectrometrists should search for all peptides, but assess only the ones they care about". *Nature Methods*, 14(7):644, 2017.

[3] R. Peckner, S. A. Meyers, J. D. Egertson, R. S. Johnson, J. G. Abelin, S. A. Carr, M. J. MacCoss, and J. D. Jaffe. Specter: linear deconvolution as a new paradigm for targeted analysis of data-independent acquisition mass spectrometry proteomics. *Nature Methods*, 15(5):371–378, 2018.

[4] P. Sinitcyn, H. Hamzeiy, F. Salinas Soto, D. Itzhak, F. McCarthy, C. Wichmann, M. Steger, U. Ohmayer, U. Distler, S. Kaspar-Schoenefeld, et al. MaxDIA enables library-based and library-free data-independent acquisition proteomics. *Nature Biotechnology*, 39(12):1563–1573, 2021.

[5] A. Sticker, L. Martens, and L. Clement. Mass spectrometrists should search for all peptides, but assess only the ones they care about. *Nature Methods*, 14(7):643–644, 2017.

[6] M. T. Strauss, I. Bludau, W.-F. Zeng, E. Voytik, C. Ammar, J. P. Schessner, R. Ilango, M. Gill, F. Meier, S. Willems, et al. AlphaPept: a modern and open framework for MS-based proteomics. *Nature Communications*, 15(1):2168, 2024.

[7] F. Yu, G. C. Teo, A. T. Kong, K. Fröhlich, G. X. Li, V. Demichev, and A. I. Nesvizhskii. Analysis of DIA proteomics data using MSFragger-DIA and FragPipe computational platform. *Nature Communications*, 14(1):4154, 2023.
